# PhyloCoalSimulations: A simulator for network multispecies coalescent models, including a new extension for the inheritance of gene flow

**DOI:** 10.1101/2023.01.11.523690

**Authors:** John Fogg, Elizabeth S. Allman, Cécile Ané

**Affiliations:** Department of Statistics, University of Wisconsin - Madison, WI, 53706, USA; Department of Mathematics and Statistics, University of Alaska - Fairbanks, AK, 99775-6660, USA; Department of Botany, University of Wisconsin - Madison, WI, 53706, USA

**Keywords:** gene tree, species network, species tree, introgression, hybridization, admixture graph

## Abstract

We consider the evolution of phylogenetic gene trees along phylogenetic species networks, according to the network multispecies coalescent process, and introduce a new network coalescent model with correlated inheritance of gene flow. This model generalizes two traditional versions of the network coalescent: with independent or common inheritance. At each reticulation, multiple lineages of a given locus are inherited from parental populations chosen at random, either independently across lineages, or with positive correlation according to a Dirichlet process. This process may account for locus-specific probabilities of inheritance, for example. We implemented the simulation of gene trees under these network coalescent models in the Julia package `PhyloCoalSimulations`, which depends on `PhyloNetworks` and its powerful network manipulation tools. Input species phylogenies can be read in extended Newick format, either in numbers of generations or in coalescent units. Simulated gene trees can be written in Newick format, and in a way that preserves information about their embedding within the species network. This embedding can be used for downstream purposes, such as to simulate species-specific processes like rate variation across species, or for other scenarios as illustrated in this note. This package should be useful for simulation studies and simulation-based inference methods. The software is available open source with documentation and a tutorial at https://github.com/cecileane/PhyloCoalSimulations.jl.

---

It is now common for genome-wide molecular data sets to be collected across many individuals from many species. Evidence for discordance between gene genealogies has been abundant for decades now (Mallet et al., 2016). For a single site in the genome or for a locus that did not experience recombination, the genealogical history of this site or locus is a tree — also called *gene tree*. It represents the history of the locus sequenced across one or more individuals from multiple species (or populations). The species phylogeny, on the other hand, represents the history of populations. It may be more complex than a tree if these populations experienced processes beyond population splitting, like populations merging or migrations of individuals between populations. Gene trees are embedded within the species phylogeny, because the individuals carrying the genes are members of the populations in the species phylogeny — but gene trees need not have topologies matching that of the species phylogeny, nor matching each other.

Beyond tree estimation error, a ubiquitous process responsible for discordance between gene trees is incomplete lineage sorting within any given population. This process is typically modelled by the coalescent (Rannala et al., 2020). When this process is applied to each population along the species phylogeny, it is referred to as the multispecies coalescent (MSC), and as the network multispecies coalescent (NMSC) when the species phylogeny is a network (Degnan, 2018). This probability model is already widely used as a basis for likelihood-based methods to estimate species phylogenetic networks (e.g. Solís-Lemus and Ané, 2016; Yu et al., 2014; Rabier et al., 2021; Allman et al., 2019a; Lutteropp et al., 2022; Blair and Ané, 2020).

Simulators are essential for many scientific endeavors, such as to assess the accuracy of inference methods, when data are simulated with known parameter values. Simulators are also used to build approximate Bayesian computation approaches and other simulation-based inference methods (Beaumont, 2010; Fan and Kubatko, 2011). We present here a simulator for the network multispecies coalescent model. It takes a species phylogenetic network as input (this network may be a tree) and simulates gene trees embedded in this network according to the NMSC. Currently, options to do this are few. The simulator `ms` (Hudson, 2002) can simulate multiple populations and migration between populations, but cannot take as input the standard extended Newick representation of a species network (Cardona et al., 2008), which makes network input prone to error. Other simulators are either restricted to species trees (e.g. SimPhy, Mallo et al., 2016) or are not reliable, like HybridLambda (Zhu et al., 2015) (see Allman et al., 2022).

In this work, we extend the network coalescent process to include correlated inheritance of lineages coexisting at a reticulation event. We implemented and describe a package to simulate gene trees under this model easily, in the fast and interactive Julia language (Bezanson et al., 2017). In the rest of this paper, we describe our model for correlated inheritance, key functionalities of our simulator, our validation of the implementation, and examples of potential downstream uses.

## 1. The network multispecies coalescent model

### 1.1. The coalescent model in one population

Within a given population, the Wright-Fisher model tracks the full set of individuals forward in time, assuming non-overlapping generations, random mating and neutrality (Hahn, 2018). The coalescent model tracks the genealogy of a sample of individuals back in time. Assuming a large haploid population size *N*, the time at which two individuals have a common ancestor is approximately exponentially distributed with rate 1*/N*, with time measured in number of generations. The process can be described more simply with time measured in *coalescent units*, which are number of generations divided by the population size. For any pair of sampled individuals, the time of their most recent common ancestor, in coalescent units, is exponentially distributed with rate 1. If the population size varies over time, the instantaneous rate of coalescence between two given individuals at a given time *t* is 1*/N* (*t*) and the distribution of their coalescent time is a generalization of the exponential distribution, described in Allman et al. (2019b).

With three or more individuals in a sample, the pair that coalesces first is uniformly distributed among all pairs.

### 1.2 Phylogenetic networks

In a species network, a speciation (or a population split) is represented by a *tree node* with one parent edge (the ancestor population) and two or more child edges (the descendant populations). A reticulation is represented by a *hybrid node* with two (or more) parent edges (the parental populations) and one child edge (the descendant population). A reticulation may arise from various biological processes, such as hybrid speciation or introgression (in eukaryotes), lateral gene transfer (in bacteria) or gene re-assortment (in viruses) for example. At a reticulation, each parent edge has an inheritance probability *γ* (or hybridization parameter), which represents the proportion of genes inherited by descendant population from that parental population. The inheritance of the partner parents need to sum up to 1. With exactly 2 parents, the parent edge with *γ >* 0.5 is called the *major* parent, and the other is called the *minor* parent.

### 1.3 The network multispecies coalescent model

The network multispecies coalescent model uses the single-population coalescent within each population, that is, within each edge in the network. At a speciation node, the gene lineages from descendant populations come together in the parent population, where they can then coalesce, going back in time. At a hybrid node, each gene lineage reaching the node (going back in time) is randomly assigned to be inherited from one of the parent populations according to their inheritance probabilities.

The typical assumption made by most network inference methods is that of independent inheritance at a hybrid node. This means that if multiple lineages of a given locus coexist at a hybrid node, then each one is assigned to a parental population *independently* of the other lineages.

Gerard et al. (2011) used a different model with *common* inheritance, in which gene lineages reaching a hybrid node are all inherited from the same parent, chosen according to the inheritance probabilities. This model has complete positive correlation, *ρ* = 1, between lineages at a given locus, and independence across loci.

### 1.4 Correlated inheritance

Independent inheritance and common inheritance are each biologically relevant under some scenarios, although independent inheritance seems more flexible. For example, consider the effect of selection, that typically differs across loci, in a scenario when population *R* receives gene flow from migrant individuals, coming from a donor population *D*. Let *A* be the admixed population after the period of gene flow. For loci resistant to gene flow, all lineages sampled from *A* might be inherited “vertically” from *R* without any contribution from gene flow. For other loci with advantageous alleles brought in by migrant individuals, most lineages sampled from the admixed population *A* might be inherited “horizontally” via gene flow, from population *D*. As the inheritance probability *γ* is taken as a global proportion across lineages and loci, variation across loci —such as that caused by selection— can result in correlated inheritance. Other scenarios may also cause a violation of the independent inheritance model.

We introduce here a model to bridge the two extremes between independent inheritance and common inheritance. To do so, we use a Dirichlet process on the set of parental populations at a given hybrid node, with base measure determined by the inheritance probabilities and concentration parameter denoted by *α*≥ 0 (Ferguson, 1973). We first describe the process in terms of locus-specific inheritance proportions. Consider a hybrid node with *k* parents and inheritance probabilities γ = (γ_1_, …, γ_*k*_). These probabilities are global: γ_*j*_ is the proportion of genes inherited from parent *i* across all lineages from all loci. We may consider variation across loci by randomly drawing γ (ℓ) = (γ_1_(ℓ), …, γ_*k*_(ℓ)) for each locus ℓ from the Dirichlet distribution 𝒟 (*α* γ_1_, …, *α* γ_*k*_), whose mean is the vector of global probabilities γ. Then, at that locus ℓ, all lineages present at the hybrid node are randomly and *independently* inherited from one of the parent populations according to γ (ℓ). The parameter *α* controls the concentration of the locus-specific proportions. If *α* is infinite, then the Dirichlet distribution 𝒟 (*α* γ_1_, …, *α* γ_*k*_) is concentrated on the single value (γ_1_, …, γ_*k*_), such that all loci share the same inheritance proportions: there is no variation across loci. As *α* gets smaller and near 0, there is more variation across loci, whereby some loci have all their lineages inherited from one parent and other loci have all their lineages inherited from another parent. Note that the locus-specific proportions γ (ℓ) are unobserved, and are re-sampled independently at separate hybrid nodes in case the network has multiple reticulations.

An alternate and equivalent definition of the Dirichlet process highlights the correlation between lineages (Ferguson, 1973). Consider multiple gene lineages reaching the hybrid node, back in time. They can be ordered in any arbitrary way. The first lineage is assigned a parent population *Y*_1_ according to the global base distribution γ. Then other lineages have an increased preference to come from the same parent as previously-assigned lineages. Namely, the parent *Y*_*n*+1_ assigned to lineage *n* + 1 is set to *Y*_*i*_ with probability 1*/*(*α*+*n*) for each *i* = 1 …, *n*, and is otherwise drawn from the base distribution γ with probability *α/*(*α*+*n*).

We show in the appendix that the correlation between the parent assignment of two lineages *Y*_*i*_ and *Y*_*j*_ is

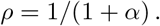

This process, therefore, bridges between the common inheritance model, which corresponds to *α* = 0, and the independent inheritance model, which corresponds to *α* = ∞.

### 1.5 Variable edge lengths and widths

Without any restrictions on the population sizes (edge widths) or the number of generations across populations (edge lengths), the model was called *modified coalescent* by Long and Kubatko (2019). In particular, the network is not required to be time-consistent, that is: multiple paths from the root to a given node may have different lengths. If the network is time-consistent then the distance from the root to a node is well-defined. The molecular clock assumption requires a network to be time consistent when edge lengths are in number of generations, and further requires the network to be ultrametric: with an equal number of generations between the root to all tips in the network. Without this clock assumption, the coalescent is called *clockless* by Long and Kubatko (2019). Note that a time-consistent network need not be ultrametric, for instance if some leaves correspond to extinct populations, or to populations that were sampled at different times.

## 2. Gene tree simulator

### 2.1 Coalescent simulation features

In `PhyloCoalSimulations`, multiple trees can be simulated at once. Trees are *independently* drawn from the NMSC, which means that they are assumed to correspond to unlinked loci. The user specifies the number of trees to be simulated, as well as the number of individuals per species. This number of individuals is constant across species if it is specified as a single number, or can vary across species if it is specified as a dictionary mapping each species name to the desired number of individuals for that species. It is also possible to simulate missing data, with 0 individuals in some species.

Our simulator implements the modified coalescent, without any restriction about the population size or the number of generations across edges in its default mode with edge lengths in coalescent units. The simulator can also take a network with edge lengths in number of generations, in which case it also needs input on the population size of each population (or edge) in the network, as well as the size of the population above the root. Given this input, each population is assumed to have a constant size over the time span of the population. The network need not be time-consistent nor ultrametric, regardless of the units (coalescent, or number of generations). In simulated gene trees, edge lengths have the same units as in the input network. The online tutorial shows how to convert these units, assuming a constant size along each population.

The user has the option to request knowledge of the embedding of gene trees within the input species network. This knowledge is provided by adding degree-2 nodes in gene trees, to map each crossing of gene tree lineages through speciation and hybridization events (see Fig. 1). In gene trees, degree-3 nodes (adjacent to 3 edges) represent coalescent events: when 2 individuals (the 2 children) coalesce into their common ancestor. Degree-2 nodes are adjacent to 2 edges only, and are created each time that a gene tree lineage crosses a node in the species phylogeny. Degree-2 nodes are named after the species node that they map to, which may be a speciation (tree) node or a reticulation (hybrid) node. When gene trees include these degree-2 nodes, each edge maps to a single edge in the species network. Therefore, one of their attributes is set to a unique identifier of the species edge that the gene tree edge maps into. The “root” population above the root of the network is given a unique identifier for that purpose. Similarly, degree-3 nodes in gene trees correspond to coalescent events that occurred within an edge in the network, and have one of their attributes track the identifier of that edge.

**Figure 1.**
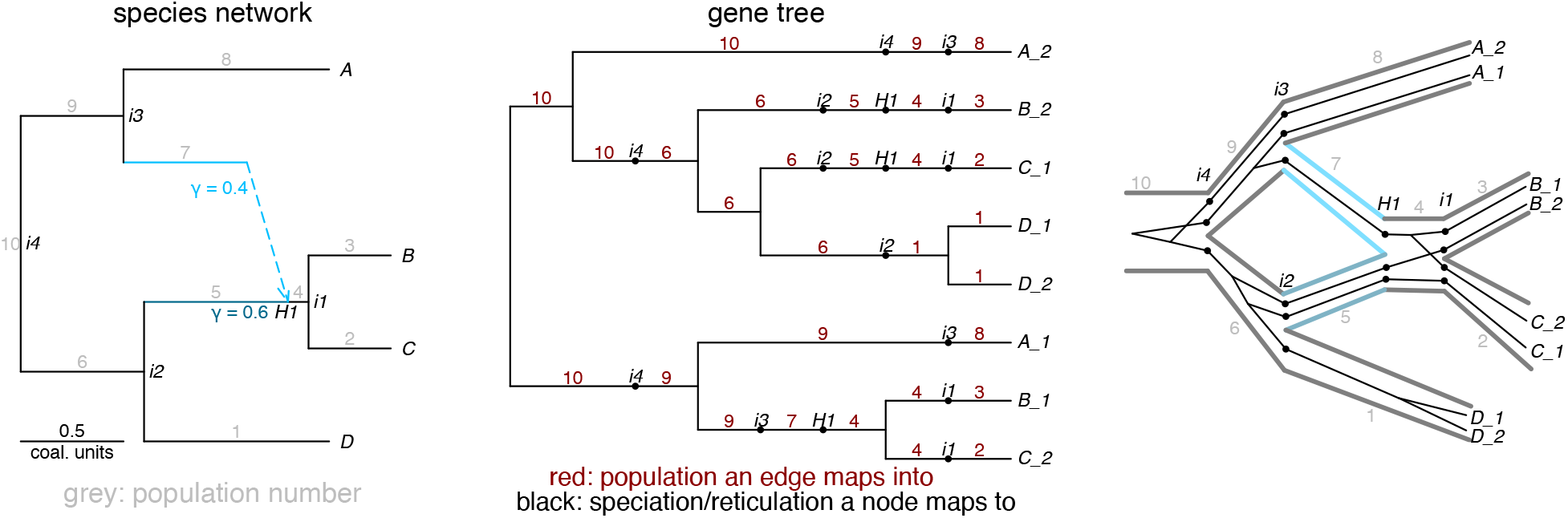
Left: example species network *N*, with edges annotated by their numbers and nodes annotated by their names. Middle: example gene tree *T* simulated from *N*, with extra degree-2 nodes (large dots) annotated by the name of the species node they map to and edges annotated by the species edge they evolved into. Right: representation of *N* with “fat” edges and *T* with thin edges embedded into *N* according to the name mapping of degree-2 nodes in *T*.

The simulated gene trees can be written to a file in Newick parenthetical format. When embedding information is requested by the user, the extra degree-2 nodes (and the length of their two incident edges) can also be written on file as part of the Newick format. As degree-2 nodes are named after nodes in the species network, the Newick description of gene trees is sufficient to store the embedding information. Other programs can then use this outside of `PhyloCoalSimulations` for downstream purposes.

### 2.2 Example use cases

We illustrate now cases that use knowledge of the embedding of gene trees within the species network. For example, one may count the number or extra lineages due to deep coalescences, following Maddison (1997). On a given edge in the species phylogeny, lineage sorting is complete if all gene lineages entering the edge (going back in time) coalesce into a single gene lineage by the time they exit the edge. If instead *k*≥ 2 gene lineages remain, then *k*− 1 of these lineages are “extra”. The total number of extra lineages, for a given gene tree, is the sum of the number of extra lineages at the end of each edge, summed over all edges in the species phylogeny. This quantity can be calculated from the known embedding of the gene tree within the network. It is of interest because inference methods may attempt to use this number of extra lineages as a criterion to infer the embedding of gene trees within a species tree or species network (Yu et al., 2013; LeMay et al., 2022; Wawerka et al., 2022).

Embedding information also allows users to count the proportion of simulated lineages inherited via gene flow. These proportions could differ from the theoretical inheritance probabilities *γ* of each hybrid edge, especially with a small number of genes (e.g., 8 loci to simulate reassortment in influenza viruses). These simulated (or realized) inheritance proportions might be of interest, for example as target quantities to estimate when assessing the performance of inference methods.

Finally, the mapping of gene tree lineages to species edges can be used to apply species-specific processes in downstream manipulations of gene trees. For example, we may scale branch lengths in gene trees to convert their original units (in number of generations or coalescent units) to a number of substitutions per site. With knowledge of the species branch that a gene lineage evolves in over time, we can apply different scaling factors to simulate molecular rate variation across species.

These illustrations are provided with example code in the package tutorial at https://cecileane.github.io/PhyloCoalSimulations.jl/dev/. The tutorial also provides example code to plot networks and gene trees, write gene trees to a file, and to suppress degree-2 nodes. These network manipulations and vizualizations are demonstrated using utilities from `PhyloNetworks` (Solís-Lemus et al., 2017, 2022) and companion package `PhyloPlots` (Ané, 2022).

### 2.3 Validation

To validate our implementation, we considered several species networks. For each network, we simulated samples of gene trees and then checked how well the distribution of select summary statistics matched the theoretical distribution under the NMSC.

#### 2.3.1 Quartet concordance factors

One summary statistic we considered focused on simulated gene tree topologies. For a given set of 4 taxa, a *quartet* is an unrooted binary topology on these 4 taxa. There are 3 distinct quartets, described by how their internal edge splits the 4 taxa into two sets of 2 taxa. On {*a, b, c, d*}, the 3 quartets are denoted as{ *ab*| *cd*}, {*ac*| *bd*} and {*ad*| *bc*}. The proportion of genes whose tree displays a particular quartet is called a *quartet concordance factor* (qCF). On a level-1 network, in which reticulations create cycles that do not share any edge, expected qCFs can be calculated in PhyloNetworks.

Since qCFs can be observed from data and their expectations can be calculated from a species network, qCFs can be used to estimate numerical parameters on a species network and infer the network topology itself as done in SNaQ (Solís-Lemus and Ané, 2016), or to estimate solely the network topology using a fast distance-based method NANUQ (Allman et al., 2019a). Quartet concordance factors are also used to test the fit of a multilocus gene tree data set to the MSC on a tree, one quartet at a time (Mitchell et al., 2019), and to test the adequacy of a candidate network under the NMSC (Cai and Ané, 2020).

We simulated 200 sets of 10, 000 gene trees, each with 1 individual per species, under the network on the left of Fig. 2, with inverse population sizes simulated from a uniform distribution between [1*/*2000, 1*/*666]. These settings were chosen to test the validity of the software when the species network parameter is input with number of generations and population sizes. In addition we simulated 2 sets of 10, 000 gene trees, each with 1 individual per species, under the middle network in Fig. 2 with branch lengths in coalescent units (with independent inheritance of lineages from *A* and *B* at the reticulation). One set used the option to create degree-2 nodes in gene trees, and the other set did not, to test the validity of the software under these two settings. The first network (with 4 leaves) had short edge lengths in coalescent units, with a high level of incomplete lineage sorting and deep-time coalescence as a primary cause of gene tree discordance. The second network (with 6 leaves) had long edge lengths in coalescent units, leading to little incomplete lineage sorting and thus hybridization is the primary cause of gene tree discordance. Together, these two networks span a wide range of expected qCFs.

**Figure 2.**
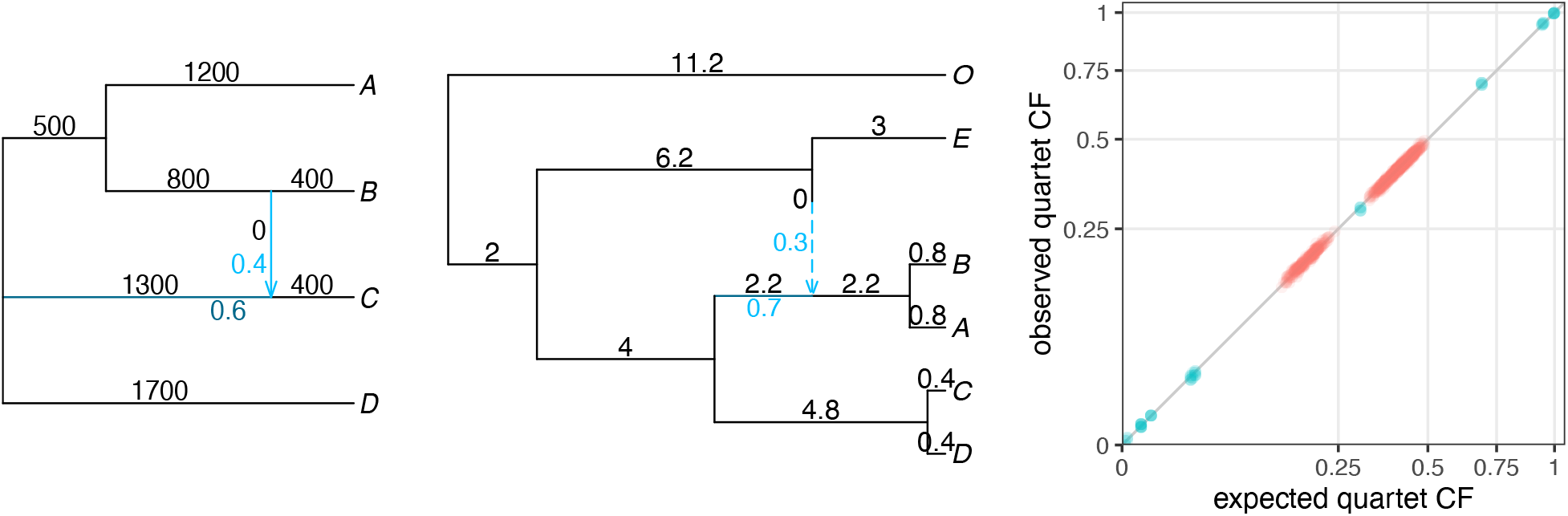
Left: network used to simulate 200 sets of 10, 000 gene trees with edges lengths in number of generations (black) and population sizes randomly selected between 666 and 2000. Middle: network used to simulate two sets of 10, 000 gene trees with edge lengths in coalescent units (black). Inheritance probabilities γ are shown in blue for hybrid edges. Right: quartet concordance factors observed in the simulated gene trees and expected from each network in red (resp. blue) from the 4-species (resp. 6-species) network, scaled with their square-root to increase resolution near 0.

From each set, we calculated the qCFs observed in the 10, 000 simulated gene trees for each set of 4 species. These observed qCFs matched the qCFs expected from the network very well (Fig. 2, right). Formal chi-square tests confirm the good fit, with p-values that averaged at 0.513, and with first and third quartiles of 0.267 and 0.744.

#### 2.3.2. Pairwise distances

For a second validation, we checked that the mean pairwise distances of a gene tree sample had the expected distribution across genes, as proposed by Allman et al. (2022), using the R package `MSCsimtester`. We first tested our implementation on the non-reticulate species phylogeny in Fig. 3 of Allman et al. (2022). This species tree has an asymmetric topology on 4 taxa, (((*A, B*), *C*), *D*), and 1000 generations between speciations. Population size *N*_*e*_ was constant over time within each branch, but varied across branches in the species tree: 2000 ancestral to *AB*, 3000 ancestral to *ABC* and 1000 ancestral to the root. Under the coalescent model on this tree, the distance between a pair of taxa has a piecewise exponential distribution. This distribution was very well matched by the empirical distribution calculated from 10, 000 gene trees simulated with `PhyloCoalSimulations` (Fig. 4), for all pairs of taxa.

**Figure 3.**
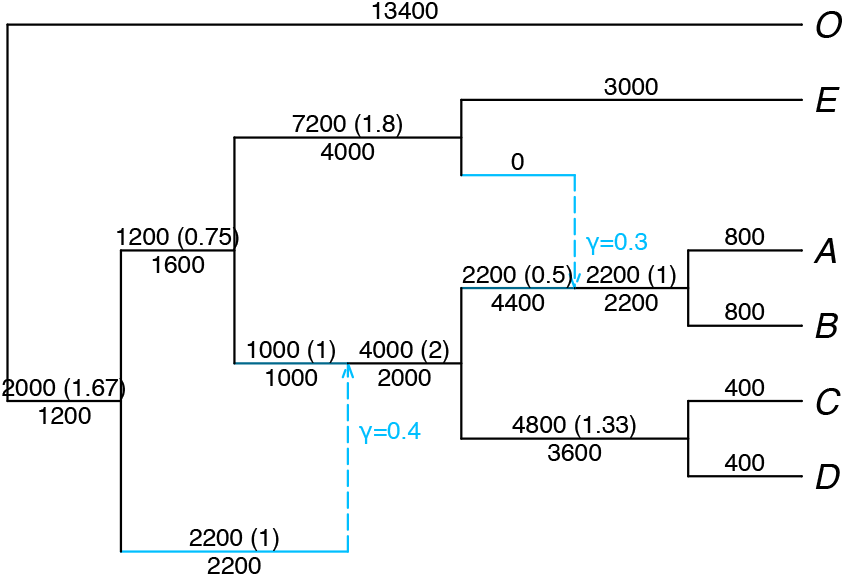
Network used to simulate 100, 000 gene trees twice (with *ρ* = 0 then *ρ* = 1) to validate the distribution of pairwise distances in gene trees. Below edges: population size *N*_*e*_. Above edges: number of generations *g* followed by coalescent units inside parentheses (*g/N*_*e*_). The population ancestral to the root had *N*_*e*_ = 1000.

**Figure 4.**
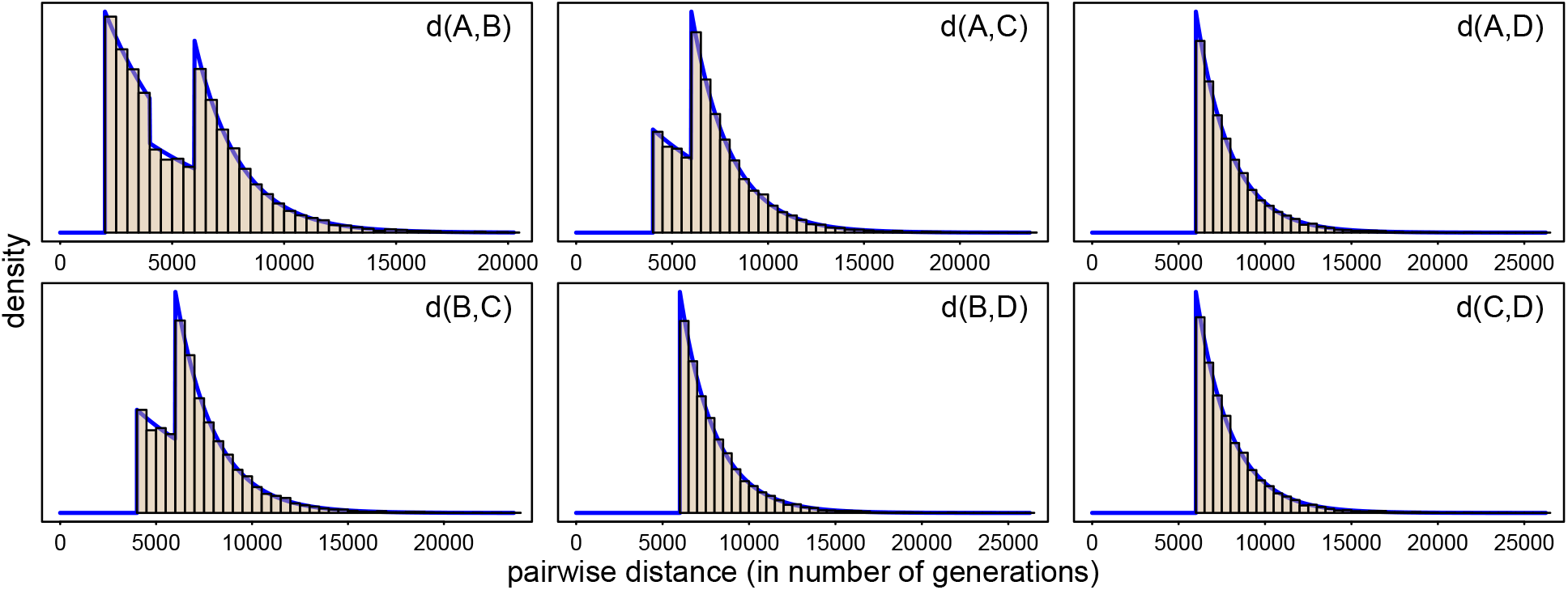
True distribution (blue curve) and distribution observed from 100, 000 simulated gene trees (brown histogram) of the distance between pairs of taxa, under the coalescent on the asymmetric tree from Fig. 3 of Allman et al. (2022): `(((A:1000,B:1000):1000,C:2000):1000,D:3000)`, and population size of 2000, 3000 and 1000 in lineages ancestral to *AB, ABC* and the root, respectively.

To validate our simulator on reticulate species phylogenies, we generated multiple gene tree samples on the network of Fig. 3, which is of level 2 as its 2 reticulations have overlapping cycles (see Huson et al. (2010) for a formal definition of level). We simulated gene trees under the fully independent model (*ρ* = 0) and the fully correlated model (*ρ* = 1). We extracted all of the network’s metric parental trees, calculated the expected pairwise distance distribution for each parental tree using `MSCsimtester`, and then for each pair of taxa combined these conditional densities using the appropriate mixture distribution to obtain the pairwise distance densities for the network. In Fig. 5, the theoretical pairwise densities are plotted together with the empirical pairwise distributions for a sample of 100,000 gene trees. Plots are shown for six of the 15 pairs of taxa, the pairs chosen to avoid redundancy from exchangeable taxon choices. Plots for all 15 pairs are in Fig. S1 in the Appendix. The match between the theoretical densities and empirical distances is visibly quite good.

**Figure 5.**
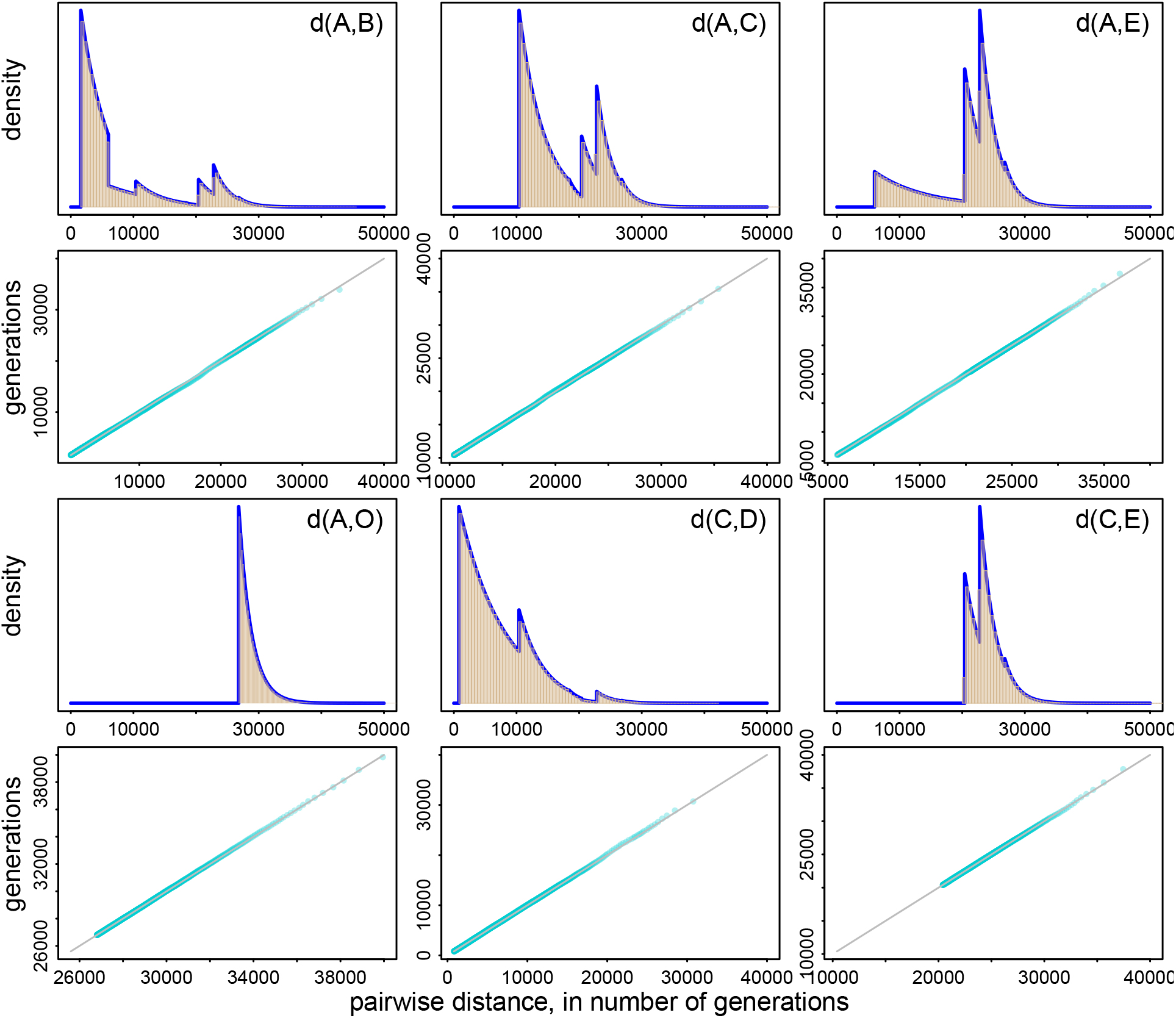
Comparison of the simulated and true distributions of pairwise distances under the coalescent (with *ρ* = 0) on the level-2 network in Fig. 3, for 6 of the 15 pairs of taxa. For each pair, the first plot (top) shows a histogram of the distances from 100, 000 simulated gene trees (brown histogram) and the true density (blue curve). The second plot (bottom, with teal points) shows a quantile-quantile plot comparing quantiles from the true distribution on the *x*-axis and quantiles from the simulator on the *y*-axis.

These plots show that the simulated gene tree distances obey the principle of ‘exchangeability’ of lineages under the NMSC. That is, if *k*≥ 2 gene lineages are present in a population simultaneously, then they are indistinguishable under the model. As a result, for example, since individuals *A, B* form a cherry on the model network, by exchangeability the empirical distance histograms for *d*(*A, X*) and *d*(*B, X*) should be roughly the same for any choice of taxon *X* different from *A, B*. This is easily confirmed by comparing plots. To quantify more precisely how good the fit of `PhyloCoalSimulations` samples was to theoretical distributions, we constructed QQ-plots for pairwise distances, also shown in Fig. 5. For these, pairwise distances were computed from samples of size 10,000 gene trees drawn from both the theoretical density (NMSC) and using our simulator (PCS). This was done for 100 replicates. For each replicate, the distances were ordered *d*_1_ ≤*d*_2_ ≤ … ≤ *d*_10,000_, and the average 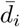 computed for each *i* = 1, …, 10000. In the QQ-plot, the pairs 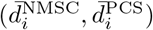 are plotted with the *x*-coordinates corresponding to theoretical values and the *y*-coordinates to the simulator sample values. The correlation is excellent.

Finally, similar validation results are in Figs. S2 and S3 for samples generated under the NSMC with the fully correlated model, *ρ* = 1, for the network of Fig. 5.

## 3. Conclusion

We extended the coalescent model on networks to include correlated inheritance of gene lineages at hybrid nodes, using a Dirichlet process to allow for intermediate correlation between 0 and 1. We implemented and validated the simulation of gene trees under the widely used network multispecies coalescent model. Our implementation in `Julia` has multiple advantages, thanks to its speed and interactivity. Once simulated, gene trees can be manipulated directly within `Julia` for downstream analyses, and can be written to a file to be used by external programs.

The embedding information carried by degree-2 nodes to map gene trees into the species phylogeny can be used in many ways, such as to extract high-level gene tree characteristics. It can also be used for customizing a study, to include the simulation of population-specific processes in downstream gene tree modifications, such as for re-scaling gene tree edges by a factor depending on the population that each gene lineage evolved in, or for the simulation of traits or molecular data whose transition rate may vary across species (or populations). Therefore, our simulation package enables the design of highly flexible simulation studies.

Further desirable functionalities of the simulator, to be considered for future versions, could include the simulation of gene duplications and losses, perhaps borrowing from the model used by `SimPhy`, which is currently limited to species trees (Mallo et al., 2016). The simulation of reticulation via polyploidization, which involves whole genome duplication, would also be very desirable. Including the effect of selection and linkage between loci are other interesting future endeavors.

## 4. Availability

The simulation package is available open source with documentation and a tutorial at https://github.com/cecileane/PhyloCoalSimulations.jl. Code to reproduce the validation studies and figures is available from the Dryad Digital Repository: http://dx.doi.org/10.5061/dryad.[NNNN].

## Acknowledgements

We thank Hector Baños, John Rhodes, and Jingcheng Xu for helpful discussions and for encouraging us to pursue this work.

## Appendix A.

### Correlation from the Dirichlet process

Consider a given hybrid node with *k* ≥ 2 parents, and let *m* be one of its *k* parental populations. We prove here that the correlation is *ρ* = 1*/*(1 + *α*) between 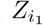 and 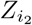, where *Z*_*i*_ is equal to 1 if lineage *i* is inherited from parental population *m* and 0 otherwise. To simplify notations, let γ = γ_*m*_ be the probability that a lineage is inherited from parent *m*. Then each *Z*_*i*_ has a Bernoulli distribution *Z*_*i*_ ∼ 𝔅 (γ) individually. Since the Dirichlet process is exchangeable, we simply need to consider the case *i*_1_ = 1 and *i*_2_ = 2. Then cor(*Z*_1_, *Z*_2_) = (𝔼(*Z*_1_*Z*_2_) − γ^2^)*/*(γ (1 − γ)) and the result follows easily from

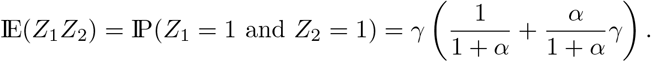

## Appendix B.

### Vpalidation of the distribution of pairwise distances

**Figure S1.**
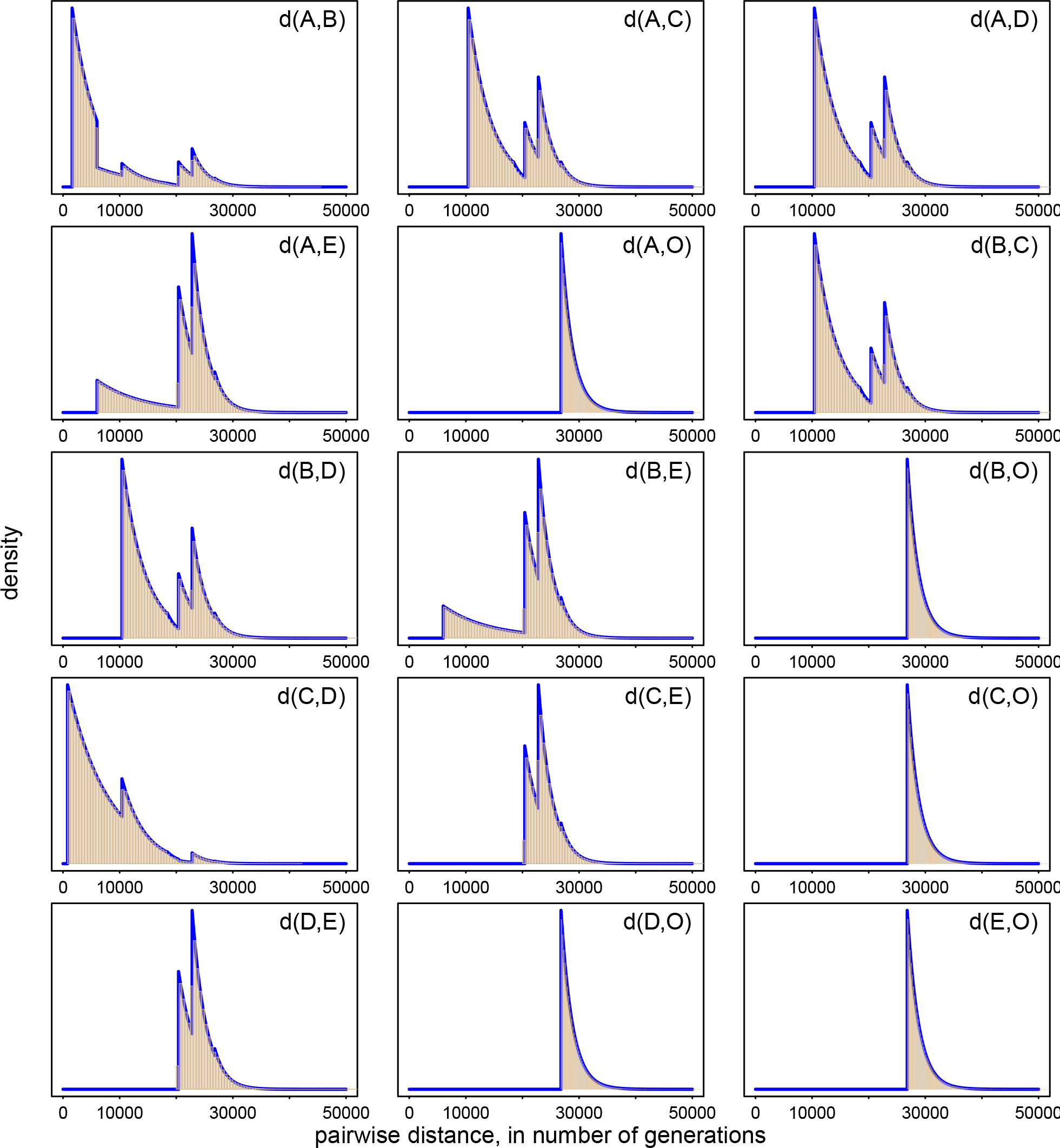
True distribution (blue curve) and distribution observed from 100, 000 simulated gene trees (brown histogram) of the distance between all 15 pairs of taxa, under the coalescent with *ρ* = 1 on the level-2 network in Fig. 3. For ease of comparison, six of the histograms in Fig. 5 are repeated here.

**Figure S2.**
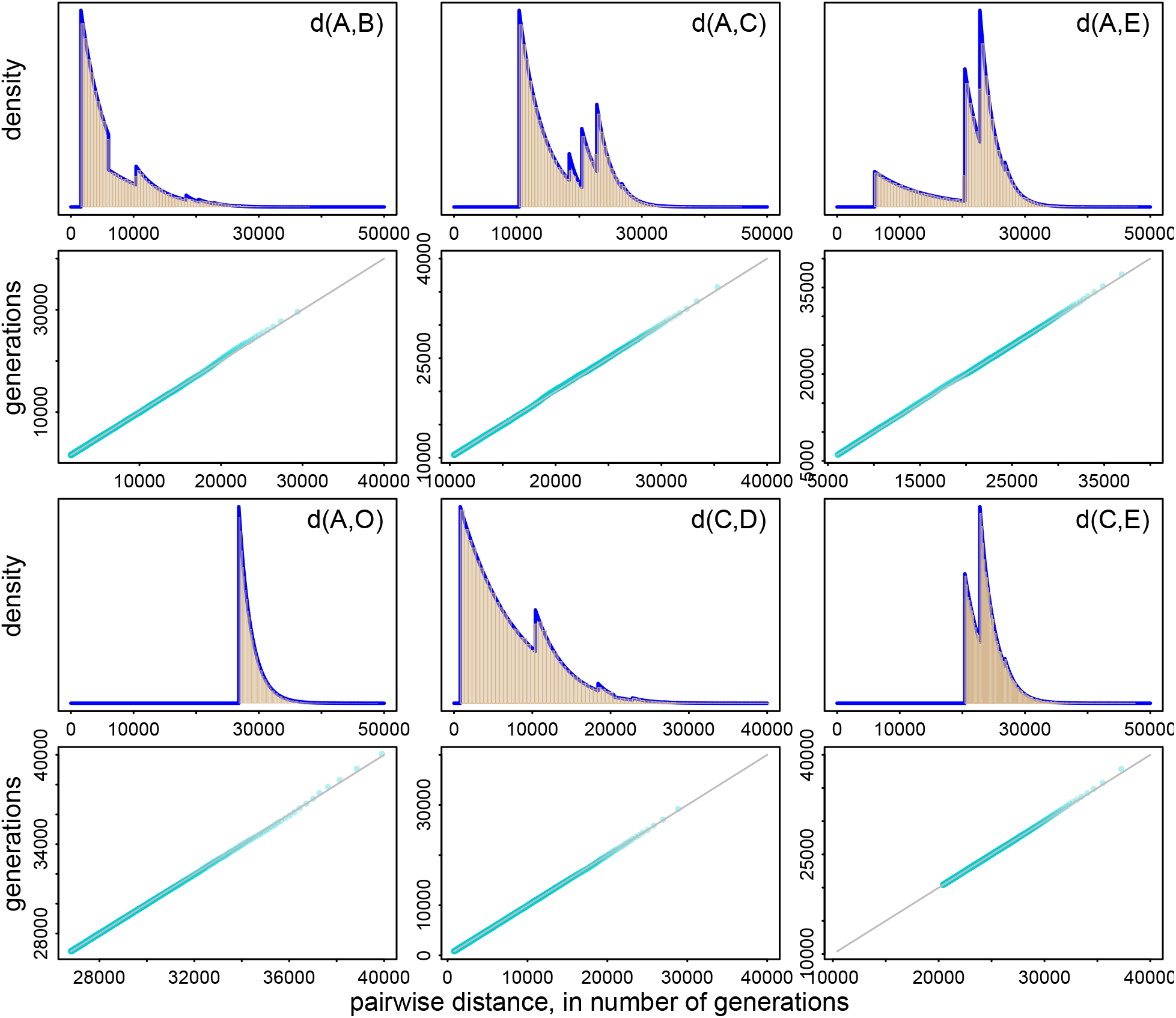
Comparison of the simulated and true distributions of pairwise distances under the coalescent on the level-2 network in Fig. 3, for 6 pairs as in Fig. 5 but with common inheritance: *ρ* = 1.

**Figure S3.**
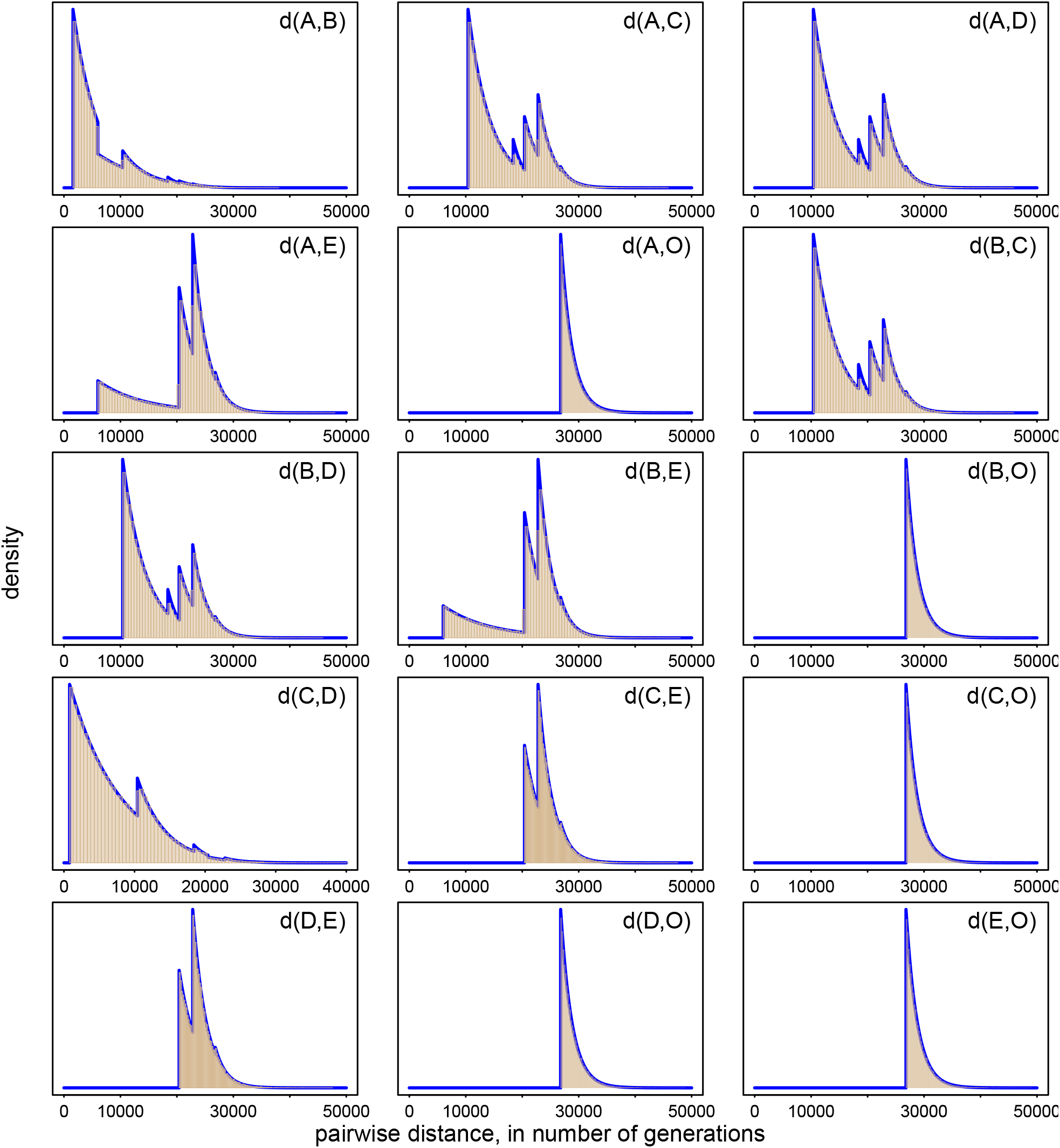
Comparison of the simulated and true distributions of pairwise distances under the coalescent on the level-2 network in Fig. 3, for all 15 pairs of taxa as in Fig. S1 but with common inheritance: *ρ* = 1.

